# phastWeb: a web interface for evolutionary conservation scoring of multiple sequence alignments using phastCons and phyloP

**DOI:** 10.1101/327569

**Authors:** Ritika Ramani, Katie Krumholz, Yifei Huang, Adam Siepel

## Abstract

The Phylogenetic Analysis with Space/Time models (PHAST) package is a widely used software package for comparative genomics that has been freely available for download since 2002. Here we introduce a web interface (phastWeb) that makes it possible to use two of the most popular programs in PHAST, phastCons and phyloP, without downloading and installing the PHAST software. This interface allows users to upload a sequence alignment and either upload a corresponding phylogeny or have one estimated from the alignment. After processing, users can visualize alignments and conservation scores as genome browser tracks, and download estimated tree models and raw scores for further analysis. Altogether, this resource makes key features of the PHAST package conveniently available to a broad audience.

**AVAILABILITY:** phastWeb is freely available on the web at http://compgen.cshl.edu/phastweb/. The website provides instructions as well as examples.

## INTRODUCTION

In recent years, there have been enormous investments in complete genome sequencing of species that fall close to one another on the tree of life, allowing for comparative genomic analyses on unprecedented scales. The PHylogenetic Analysis with Space/Time models (PHAST) software package has emerged as a popular and widely used toolkit for analyzing such comparative genomic data. PHAST is best known as the engine behind the Conservation tracks in the University of California, Santa Cruz (UCSC) Genome Browser, but it additionally includes several programs for phylogenetic modeling and functional element identification, as well as utilities for manipulating alignments, trees and genomic annotations.

Since 2002, PHAST has been available as a collection of command-line programs and supporting software libraries that users must download and install to apply to their own sequence data. However, traffic on the PHAST mailing list indicates that many users are exclusively interested in producing conservation scores or predicted conserved elements using the phastCons or phyloP programs. This application of PHAST is fairly straightforward and only requires a handful of programs. In addition, users often wish to visualize their conservation scores and predicted conserved elements together with their multiple sequence alignment in a Genome Browser display, but the existing PHAST package does not support such visualization.

Here we introduce an easy-to-use web interface to PHAST, called phastWeb, to facilitate conservation-scoring using PHAST and visualization using the UCSC Genome Browser. Users of phastWeb are able to circumvent the non-trivial process of installing the PHAST software, running several command-line tools, converting output formats, and uploading data for visualization. Instead, all of the necessary steps are launched via a self-explanatory user interface and executed on our servers. Visualization is accomplished using the UCSC Genome Browser’s “track hub” mechanism (Raney et. al., 2014). Users of phastWeb can either estimate phylogenetic trees, branch lengths, and substitution models from their own data sets, or accept pre-estimated models. Key intermediate data files (such as phylogenetic models and “wig” files of conservation scores) are made available for download.

## METHODS

### Getting started

The only required input for phastWeb is a sequence alignment file (in MAF, FASTA, or PHYLIP format). In addition, the user may optionally provide a known phylogeny or pre-estimated neutral model (*.mod) file, if one is available from a previous analysis. All computations are accomplished on the server side, using phastCons, phyloP, and other programs from PHAST, together with the phastWeb scripts.

### Estimating the neutral model [if not provided]

If the user has not provided a *.mod file, a neutral model must be estimated from the alignment. This step requires a tree topology defining the phylogenetic relationships of the aligned sequences. The user can choose to upload a known tree topology, such as a published tree or one that has been estimated separately, or have the topology estimated from the alignment using the neighbor joining method. If necessary, tree estimation is accomplished using the *neighbor* program from PHYLIP (Felsenstein, 2005). Once a tree is obtained, a neutral substitution model is estimated from the data using the phyloFit program in PHAST. The user has the option to upload a file defining the locations of sites likely to be free from the influence of natural selection (such as fourfold degenerate sites in coding regions, ancestral repeats, or intergenic regions). If this option is not selected, the model is estimated from all sites in the alignment under the assumption that most sites are not under selection (as is typically true for large genomes). The user also can select from one of several nucleotide substitution models implemented in PHAST.

### Running phastCons and phyloP

Once a neutral model is obtained, the user can proceed with conservation scoring. The user has the option to run the phastCons and phyloP programs independently or together. Sensible default parameters are provided for both programs but the user is free to customize them as desired, with guidelines provided in the online instructions.

### Output

When a job is submitted, phastWeb provides an estimate of the required run time based on the size of the alignment and the number of species. Once the results are available, users receive an email with a link to a results page presented in three main parts, including: (1) a link to the UCSC Genome Browser’s track hub displaying the generated conservation scores together with the reference genome and alignment; (2) a zip file containing the phastCons and/or phyloP results (*.wig files), the tree topology (if estimated by *neighbor*), the neutral phylogenetic model estimated by phyloFit, and the bigwig files for UCSC Genome Browser; and (3) an image (in scalable vector graphics [svg] format) displaying the neutral phylogeny used for the analysis.

## CONCLUSION

phastWeb is an easy-to-use web-based interface to PHAST designed for users who wish to produce conservation scores or predict conserved elements from their own multiple sequence alignments, without downloading and installing the PHAST package. The system allows visualization of predictions in the UCSC Genome Browser and estimation of phylogenies and neutral models as needed. In addition to providing the basic functionality needed by many users, phastWeb can serve as an entry-point to a more elaborate conservation analysis using the full PHAST toolkit. If the phastWeb interface proves sufficiently useful, we may extend it to include other programs in PHAST, such as phyloFit and phastBias.

**Figure 1.**
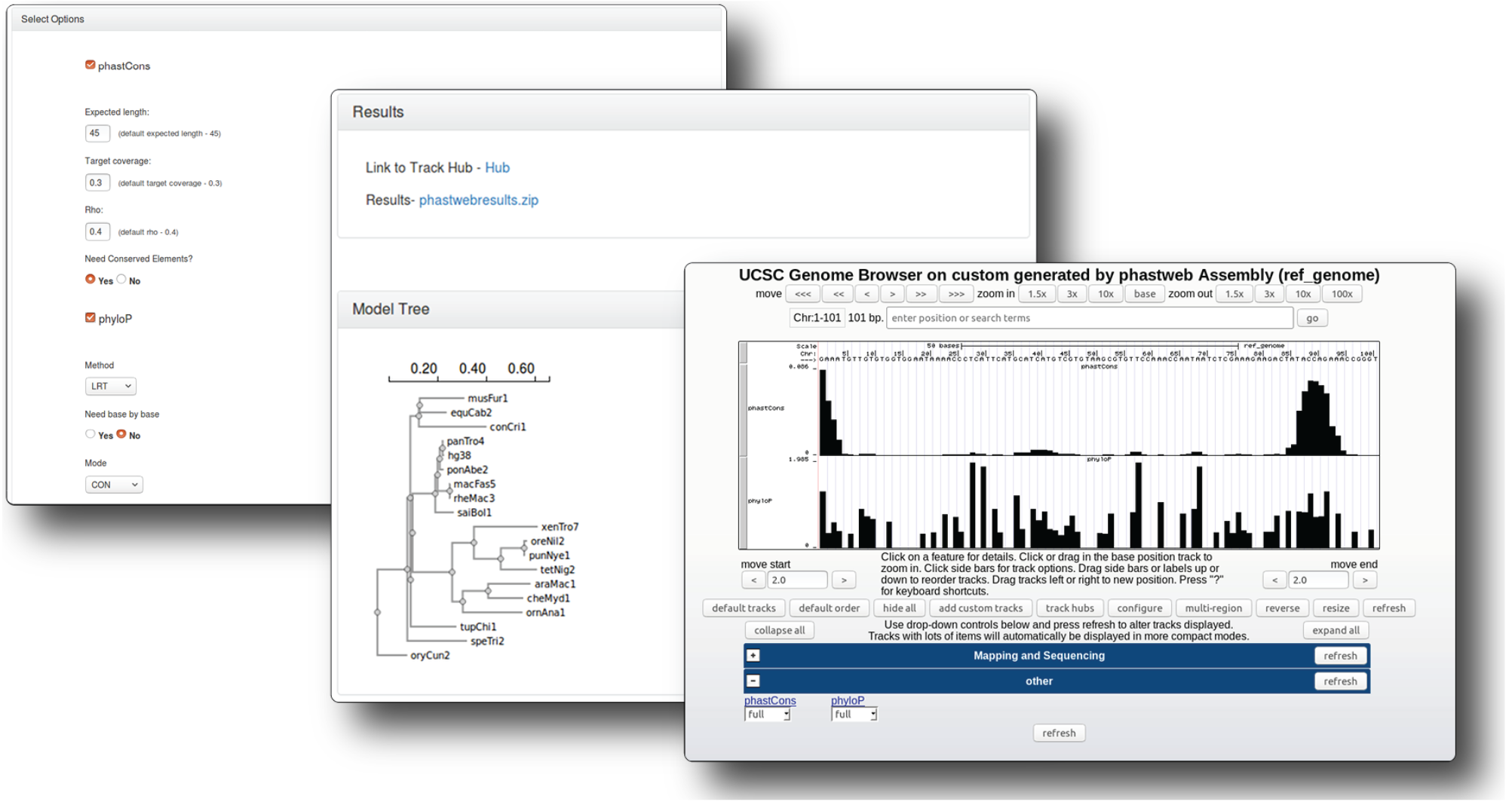
Web Interface for phastWeb. The interface prompts users to select options for running phastCons and phyloP programs independently or together after a .mod file is provided or a neutral model has been estimated from the alignment. The results page presents the zip file of phastWeb result with tree topology and a link to the UCSC Genome Browser’s track hub displaying generated conservation scores together with the reference genome and alignment.

## ACKNOWLEDGMENTS

We thank Noah Dukler for preparing Figure 1 and designing the logo for the phastWeb website.

